# Genomic Characterization of 30 Lytic *Klebsiella pneumoniae* Bacteriophages

**DOI:** 10.64898/2026.01.22.701196

**Authors:** Daniel Mora, Trisha Gryder, Jordyn Michalik-Provasek, Michael Satlin, Thomas J. Walsh, Jason J. Gill

## Abstract

The spread and rise of antimicrobial resistance poses a risk to public health due to limited effective treatment options. Alternative antimicrobials that are effective against gram-negative multi-drug resistant pathogens. The increasing rate of carbapenem resistance observed in *Klebsiella pneumoniae*, indicates the need for alternative antimicrobial options. Bacteriophages that target Klebsiella pneumoniae are promising alternative antimicrobial option, with successful treatments being reported. Here we characterized 30 lytic bacteriophages from various environmental sources and tested their effectiveness against nine clinically relevant carbapenem-resistant *K. pneumoniae* isolates. These phages were characterized through genomic sequencing, bioinformatic analysis, virulence in liquid medium, and host range on different mediums. Bioinformatic analysis revealed a diverse collection of phages that span 9 ICTV recognized families and 13 genera with genome sizes ranging from 39-349 kbp. The phages were able to inhibit bacterial growth, and no virulence or antibiotic resistance genes were detected within the phage genomes. Host range testing demonstrated phages with broad host range have varying infectivity when plated on different common growth mediums. This study includes candidate phages for further potential development as potential antimicrobial agents against CR-KP, and the complexity in understanding phage-host dynamics of non-capsule phages that target against *K. pneumoniae*.

## 1. Introduction

*K. pneumoniae* is a gram negative, non-motile, rod-shaped bacterium in the Enterobacteriacea*e* family. It is known for its polysaccharide capsule, antibiotic resistance and ability to be both a human commensal and opportunistic pathogen [1,2]. *K. pneumoniae* can colonize human mucosal surfaces and skin asymptomatically and is capable of causing a range of serious infections including pneumonia, liver abscesses, urinary tract infections, and sepsis [1]. *K. pneumoniae* can be acquired from healthcare or community settings, and infections have become difficult to treat due to its resistance to frontline antibiotics [1,3]. *K. pneumoniae* is part of a group of human pathogens commonly associated with high level of antibiotic resistance designated the ESKAPE pathogens that includes *Enterococcus faecium, Staphylococcus aureus, Klebsiella pneumoniae, Acinetobacter baumannii, Pseudomonas aeruginosa* and *Enterobacter* spp. [4].

The rise of antibiotic resistance in *K. pneumoniae* is a threat to public health globally [2,4]. In 2022, it was estimated that 4.71 million deaths were associated with bacterial antimicrobial resistance, and included 1.14 millions deaths that were attributed to bacterial drug resistance [5]. The WHO has recognized carbapenem resistant *K. pneumoniae* (CR-KP) as pathogen that poses the most urgent threat to public health [2]. The classification of CR-KP as a microorganism of concern is due to the high trend of antibiotic resistance, low preventability, and high mortality outcomes due to limited treatment options [2,6]. This is due to the fact that CR-KP harbor extended-spectrum-beta-lactamase (ESBL) enzymes, which make them resistant to nearly all beta-lactam antibiotics [7–9]. The most notable enzymes are the *bla*_*KPC*_ alleles, which were first identified in the United States in 1996 and can hydrolyze a wide range of beta-lactams [10,11]. CR-KP has been implicated as the driver for the rising increase of carbapenem resistance globally [12,13]. The spread of drug resistance has led to an urgent need for antimicrobials effective against drug resistant pathogens [2,5].

A promising alternative to traditional antibiotics is phage therapy, the use of phages as antimicrobial agents against pathogenic bacteria. Phages were first discovered in the early 20^th^ century by Felix d’Herelle, who isolated an agent effective against Shiga dysentery [14]. D’herelle coined the term bacteriophage “bacteria-eater” and pioneered the early use of phage therapy [14,15]. Phage therapy was largely abandoned in the 1940s following the introduction of antibiotics [15,16]. With the rise of antibiotic resistance and challenges developing novel and effective antibiotics, this has prompted renewed interest in phage therapy [17,18].

There are several factors that make phages an attractive alternative antimicrobial agent. Phages are ubiquitous in nature, estimated at 10 ^31^ phages particles in the biosphere, can be isolated readily isolated from the environment [19–21]. Additionally, phages are highly specific to their bacterial hosts, and can typically only infect certain strains of their host [22]. This specificity is largely due to the tail fibers or tail spikes of phages only recognizing specific receptors on the host cell surface [23]. This host specificity means that phages can target pathogens without disturbing the other members of microflora [24]. These properties and demonstrated efficacy in reducing the bacterial burden *in vitro* and *in vivo* have renewed interest in the application of phage therapy [25–29]

There are over 1600 phages able to infect *Klebsiella* hosts listed on NCBI virus [30]. Phages that target *K. pneumoniae* have been isolated from a wide range of sources, including soil, water, sewage, and human feces [22,31]. Several studies have revealed a large genomic diversity of phages that can target *Klebsiella* spp. [22,31]. For the successful application of phage therapy, careful characterization of environmental phages are necessary. Phages must be analyzed to ensure they follow a strictly lytic lifecycle and do not carry any virulence or antibiotic resistance determinants [32]. In addition, determination of phage host range and efficacy in reducing bacterial load *in vitro* against clinically relevant strains are commonly performed to identify suitable therapeutic phage candidates.

To address the urgent need for new antimicrobials against CR-KP, this chapter focuses on the isolation and characterization of 30 lytic phages able to infect clinically relevant *Klebsiella* spp. strains [33]. These phages were isolated from diverse environmental sources; influent wastewater, activated sludge, surface waters, and swine feces. The phages include a variety of morphologies spanning nine ICTV families and 13 distinct genera [34,35]. We also measured each phage’s infectivity on different clinically isolated *Klebsiella* spp. and under various culture conditions to define their host range. Together the genomic and experimental data reveals each phage’s taxonomic identity and its potential for future use. The goal of this study is to present a comprehensive overview of these 30 *Klebsiella* spp. phages to lay the groundwork for selecting the most promising candidates as antibacterial agents against carbapenem resistant *Klebsiella* spp.

## 2. Materials and Methods

### 2.2.1 Bacterial Strains and Culture Conditions

The *Klebsiella* strains used in this work are listed in Table 1. All culturing methods were carried out in either Luria Broth (LB) (10 g/L tryptone (BD), 10 g/L NaCl, and 5g/L yeast extract (BD)), or tryptic soy broth (TSB) (Bacto TSB, BD), with aeration at 37 °C. Solid media (LB agar or tryptic soy agar (TSA) were composed of the broth media amended with 15 g/L Bacto agar (BD). The Klebsiella clinical isolates were provided by Karen Frank of the NIH Clinical Center, or by Dr. Micheal Satlin of Weill Cornell Medical College as denoted in Table 1.

**Table 1.** Panel of *Klebsiella spp*. with bioinformatically determined Sequence Type, Capsule Type, and LPS Type.

| Species | Strain | Strain Type | Capsule (K) Lo-<br>cus Type | LPS O-antigen (O) Locus Type | # of Phages Infect-<br>ing Strain | Source |
| --- | --- | --- | --- | --- | --- | --- |
| *Klebsiella pneumoniae* | 39427 | 258 | 107 | O1/O2v2 | 9 | M. Satlin, WCMC |
| *Klebsiella pneumoniae* | 39428 | 258 | 106 | O1/O2v2 | 8 | M. Satlin, WCMC |
| *Klebsiella pneumoniae* | 43421 | 17 | 25 | O5 | 5 | M. Satlin, WCMC |
| *Klebsiella pneumoniae* | 39904 | 3585 | 111 | O3b | 1 | M. Satlin, WCMC |
| *Klebsiella quasipneumoniae* | I0201559 | 3861 | 142 | O3/O3a | 2 | K. Frank, NIH CC |
| *Klebsiella pneumoniae* | 42195 | 3586 | 102 | O1/O2v1 | 2 | M. Satlin, WCMC |
| *Klebsiella quasipneumoniae* | I0201547 | 359 | 51 | O1/O2v1 | 2 | K. Frank, NIH CC |
| *Klebsiella pneumoniae* | 44818 | 35 | 22 | O1/O2v1 | 3 | M. Satlin, WCMC |
| *Klebsiella pneumoniae* | 44809 | 1502 | 125 | O5 | 1 | M. Satlin, WCMC |

### 2.2.2 Isolation and Culture of Bacteriophages

Phage were isolated over a seven year period (2012-2019) from wastewater samples, listed in Table 2. Phages were isolated by either direct plating filter-sterilized (0.22 µm, PES membrane) wastewater samples to host lawns, or by a culture enrichment method, using 1-10 mL of filter-sterilized sample added to actively growing host cultures in TSB and incubated overnight at 37 °C with aeration [36]. Each phage was plaque purified three times to ensure clonality. Phage were routinely cultured and enumerated by the soft agar overlay method using TSA bottom plates overlaid with lawns of 5 mL top agar (15 g/L Bacto TSB, 4 g/L Bacto agar) inoculated with 100 µL of fresh overnight host culture, and incubated overnight at 37 °C [37]. Phage plaque morphology was assessed by observing and photographing phage plaques as they appeared on soft overlay lawns. High-titer stocks were prepared by inoculating log-phase cultures with phage at an multiplicity of infection (MOI) of ∼0.1 and incubating with aeration at 37 °C. Phage lysates were then centrifuged (10,000 x g, 4 °C, 10 min), filter-sterilized, by passage through 0.2 µm PES filters (Millipore). Phages were routinely diluted in phage buffer (5.8 g/L NaCl, 1.0 g/L MgSO4·7H2O, and 25 mM Tris-HCl pH 7.4).

**Table 2.**
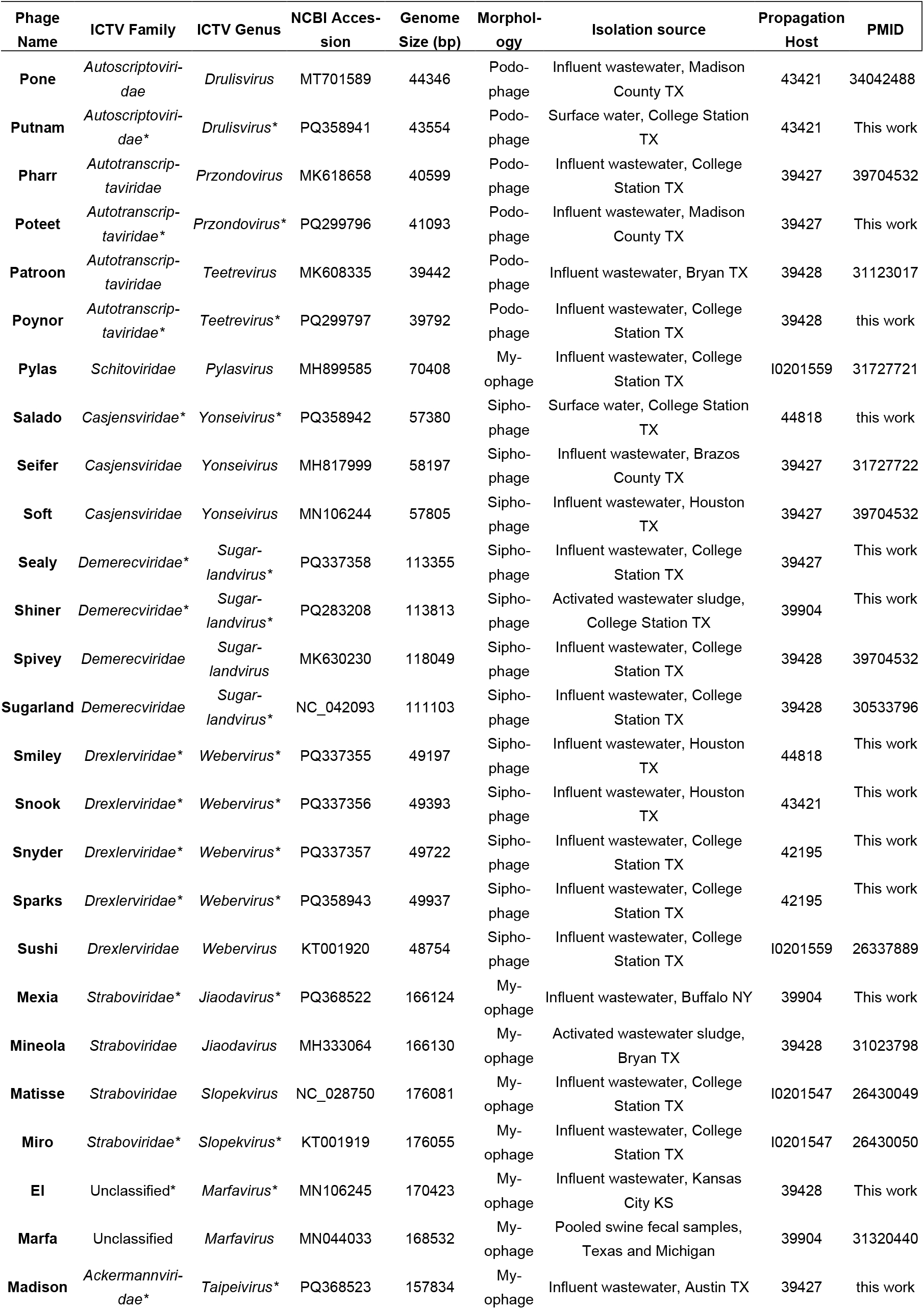
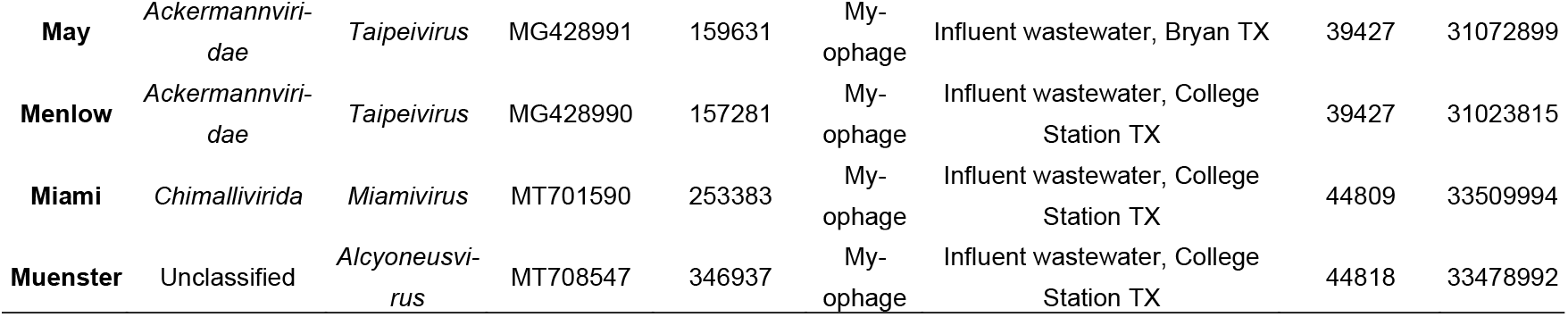
Bacteriophages described in this study. Thirty bacteriophages infecting clinical K. pneumoniae were isolated from 2012-2018 from various sources, predominantly Texas municipal wastewater. Phage taxonomic classifications at the Family and Genus levels are provided based on the current ICTV taxonomy (2024 release). Morphology is predicted based on taxonomic classification. * Represent unknown ICTV classification and use of BLAST to determine closest relative in ICTV.

### 2.2.3 DNA extraction and sequencing

Phage genomic DNA was purified from lysates using a modified protocol from the Promega Wizard DNA purification kit as described previously [38,39]. Bacterial DNA was prepared from fresh overnight cultures using the EZDNA Bacterial DNA kit (Omega Bio-Tek) kit following manufacturer instructions. Phage genomic DNA was sequenced by 454 pyrosequencing (phages Poteet and Shiner) or Illumina (all others) at commercial sequencing providers. Bacterial DNA was sequenced by Illumina at a commercial provider.

### 2.2.4 Bioinformatics

Phage genomes sequenced by 454 were assembled with Newbler v2.7. Phages sequenced by Illumina were quality controlled with FASTQC and assembled using SPAdes v3.5.0 [40,41]. Phage genomes were closed by resolution of circular assemblies or by PCR followed by Sanger sequencing. Genomes were annotated using the Center for Phage Technology’s Galaxy-Apollo phage genome annotation platform [42]. Bacterial genomes were assembled using SPAdes v3.5.0 and annotated using Prokka [40,43]. Kaptive Web was used to determine the capsule (K) and LPS (O) antigen locus types of the 9 *Klebsiella* isolates used as phage hosts [44–46]. PathogenWatch [47] was used to determine the clonal groups and Kleborate [48], was used to determine virulence and antimicrobial resistance determinants. CARD [49] was used to determine antibiotic resistance determinants, VFDB [50] was used to determine virulence determinants, and PHASTEST [51] was used to determine lysogeny determinants.

### 2.2.5 Host Range Testing

Phage host range was determined by spot testing using serially diluted phage lysates standardized to an initial titer of 10^8^ PFU/ml. Each standardized phage lysate was used to generate tenfold serial dilutions and 10 µL spot of these dilutions were plated onto a bacterial lawn, prepared as described above [37]. Phage plaques were enumerated and the apparent titer was divided by the titer of observed on the phage propagation host to calculate phage efficiency of plating (EOP) for each test strain.

### 2.2.6 Virulence Assay

Phage virulence was characterized against the propagation host using a microtiter plate-based liquid virulence assay [52]. The bacterial inoculum was prepared by subculturing bacterial overnight cultures in 1:100 in TSB and incubating with aeration at 37°C to an OD_550nm_ of 0.25 (∼10^8^ CFU/ml). Phage lysates were titered adjusted to concentration of 1×10^8^ PFU/ml. For each assay, 180 µL of adjusted bacterial inoculum in TSB was mixed with 20 µL of phage in sterile, untreated Falcon (Corning) 96-well transparent plates to achieve an initial MOI of 0.1. The plates were incubated at 37°C with double orbital shaking in a Tecan Spark 10 M plate reader (Tecan Group Ltd., Männedorf, Switzerland) and growth was monitored by measuring OD_550nm_ at 15 minute intervals for 20 hours. All assays were performed in triplicate biological replicates. Resulting growth curves were visualized in GraphPad Prism after adjusting readings against the baseline medium-only control [53].

## 3. Results

### 3.1 Characteristics of Carbapenem Resistant Klebsiella spp. Clincal Isolates

A total of nine isolates of *Klebsiella* spp. were used in this study, as shown in Table 1. Multilocus sequence typing (MLST) analysis revealed that the nine isolates were distributed across eight different strain types [47]. Two strains were identified as ST258, which has been implicated in the global spread of *K. pneumoniae* carbapenemases [54]. The remaining isolates were classified into individual ST’s: ST17, ST3585, ST3586, ST3861, ST359, ST35, ST1502. The capsule (K) and lipopolysaccharide O-antigen (O) loci were predicted bioinformatically using Kaptive, as shown in Table 1 [45]. All isolates had unique predicted K-loci, whereas only three distinct O-loci were identified. The predominant O-locus type is O1/O2, with two being predicted to have subtype O1/O2v1 and two were subtype O1/O2v2 [48]. Four isolates were predicted to have O5 locus type, and two isolates were predicted to have the O3 locus type. The remaining isolates were predicted to have O3b or O5 [48].

All isolates were found to carry resistance determinants against penicillin [8]. The one isolate did not carry a carbapenemase, the antibiotic resistant determinants found were SHV-11, CTX-M-14, and DHA-1. SHV-11 is a narrow spectrum beta-lactamase, and two ESBLs CTX-M-13 and DHA-1 [55–57]. DHA-1 is a plasmid mediated gene regulated by the ampR gene, and has been shown to be induced by carbapenems [58]. This induction leading to increased levels of DHA-1, which can confer carbapenem resistance in non-carbapenemase producing organism without the presence of OmpK36/OmpK35, which were not carried in the isolate [59,60].

Since bacterial capsule has been shown to be a limiting factor for *Klebsiella* phages, each isolate was analyzed to determine if they produced capsule [22]. The core genes that comprise the capsule of *Klebsiella* (K) locus are *galF, cpsACP, wzi, wza, wzb, wzc, wcaj/wbaP, gnd*, and *ugd*, which encode the core capsule biosynthesis machinery [61,62]. Analysis using Kaptive to confirm the that eight of the nine strains contain the core capsule locus [45]. A single isolate, *K. pneumoniae* strain 39428, was confirmed to have a nonfunctional capsule locus due to the truncation of a *wcaJ* gene.

### 3.2 Isolation, Genomic Characterization, and Diversity of Phages

The 30 isolated phages capable of infecting the nine clinically isolated *Klebsiella* spp. (Table 1) were obtained from influent wastewater, activated sludge, surface waters, and swine feces around TX and isolated using standard isolation techniques [36]. These phages were generally named with the first letter associated with the morphotype (“M” for myophages, “S” for siphophages and “P” for podophages) and named based on selected towns in Texas; the only exception to this system was the myophage EI. Examination of plaque morphology revealed that of the 30 pahges isolated, fifteen phages formed diffuse halos (Figure 1).

**Figure 1.**
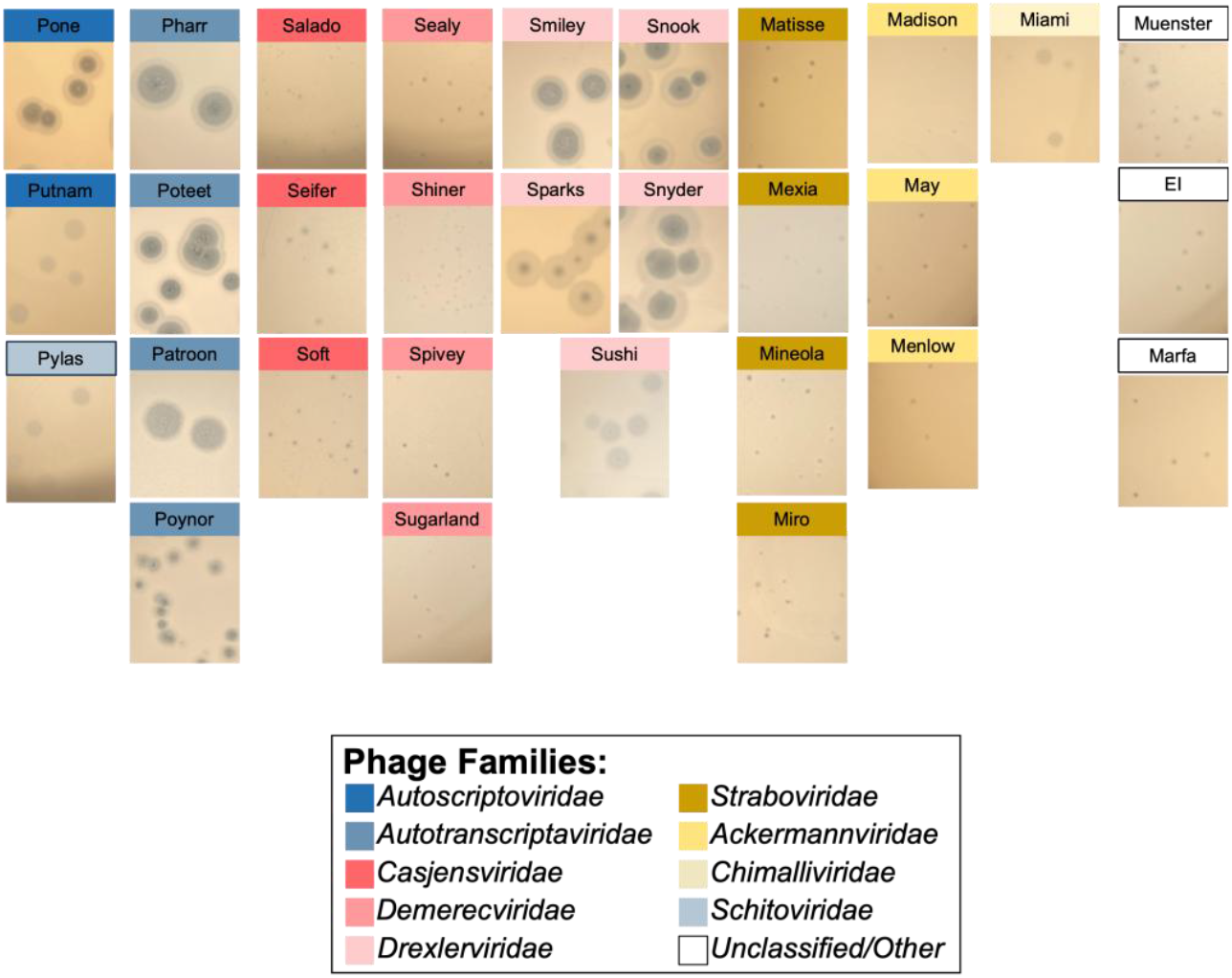
Plaque morphology of phage isolates. Phage names are colored based on ICTV taxonomic Family. All Autoscriptoviridae, Autotranscriptaviridae, Casjensviridae, Drexlerviridae, Schitoviridae, Chimallivirida, and Muenster produce haloed plaques, consistent with the presence of a predicted capsular depolymerase tail fiber or spike. Phages Patroon and Poynor lack enzymatic tail spikes but their plaques are still surrounded by halo-like zones. Casjensviridae group phages produce small haloes despite the presence of a predicted depolymerase-like tail spike. This indicates that haloed plaque formation is not a reliable indicator of phage capsular depolymerase presence.

Phage genomic DNA was purified from high titer stocks, resulting in 30 novel, nonredundant phages cultured on eight *Klebsiella* spp. isolates (Table 2). The resulting genomes ranged in size from 39,442 to 346,937 bp in length with G+C contents between 31.86% to 56.45%. Phage morphotype was predicted based on phage phylogenetic relationships and the presence of genes encoding tail components. Genome size comparisons based on morphology showed that the 11 myophages had an average genome size of 190,764 bp, the 12 siphophages had an average genome size of 73,058 bp, and the 7 podophages had an average genome size of 45,604 bp.

The phages were classified into a family and genus based on the current ICTV taxonomy (2024) release. Of the 30 phages, 15 were already placed into the ICTV taxonomy (2024). To identify an ICTV taxa for the remaining phages, the unclassified phages were analyzed by BLASTn [63] to identify the closes relative with an ICTV taxonomic designation, which was used to determine phage family and genus (Table 2) [64]. The unclassified phages showed less than 95%, but more than 70% genome similarity were vB_KleM-RaK2 (Muenster), Klebsiella phage vB_KpnP_SU552A (Putnam), Klebsiella phage KPN N98 (Salado), Klebsiella phage ABTNL-2 (Snyder), and Klebsiella phage vB_KpnS-VAC5 (Sparks). The remaining phages displayed over 95%, but less than 100% genome similarity Klebsiella phage Menlow (Madison), Klebsiella phage Pharr (Poteet), Klebsiella phage 6996 (Poynor), Klebsiella phage vB_KpnS_Uniso31 (Sealy), Klebsiella phage vB_Kpn_B01 (Shiner), Klebsiella phage GZ9 (Smiley), Klebsiella phage vB_KpnD_PeteCarol (Snook), Klebsiella phage vB_KaeM_Nis-pero (Mexia), and Klebsiella phage KP15 (Miro), and Klebsiella phage Marfa (EI).

The genomes of the phages encode between 47 and 567 genes. No virulence determinants, antimicrobial 218 determinants, and known lysogeny genes were found in any phage genomes, indication the collection is com-219 posed entirely of virulent phages [49–51]. Functional genes were divided into seven groups based on putative 220 function; DNA metabolism, head structure, lysis, tail fiber/tail structure, DNA packaging, hypothetical function 221 (Figure 2). A proteomic tree was constructed using ViPTree to show global relationships of the phages (Figure 222 3) [65]. The proteomic tree was able to classify Muenster with other phages, Klebsiella phage vB_KleM_RaK2 223 and Klebsiella phage K64-1, which were classified in ICTV under the *Alcyoneusvirus* genus.

### 3.3 Lytic Potential and Host Range of Phages

The lytic spectrum of the 30 phages were determined using nine clinically isolated carbapenem resistant *Klebsiella* spp. isolates (Figure 4, 5). A lysis curve of each phage with their respective propagation host in TSB at an MOI of 0.1 (Figure 4). Half of the phages in the collection were isolated against 39427 or 39428, whereas strain 44809 has only a single phage propagated on it. This contrasts with most strains, which have two or three phages isolated against them. The distribution of isolated phages is similar to that of *Klebsiella* spp. with common K-loci, where K types 107 and 106 represent the first and third most prevalent K-loci, respectively [48].

All phages demonstrated ability to control bacterial growth in liquid culture, typically showing an initial reduction in culture by a rebound in bacterial growth after four to eight hours of growth. Generally, the lysis curves of the phages that were propagated on the same strain had similar lysis curves. Several phages such as Sparks, Snyder, Sushi, Miro and Matisse showed an initial reduction in culture density that was followed by a rebound of bacterial growth. Following the rebound of bacterial growth there was a significant drop in culture density, that was then followed by a subsequent rebound. The myophages of 39428 EI, Mexia, and Mineola showed slightly improved ability to control bacterial growth compared to the podophages and siphophages.

Plaque assays were conducted in both TSB and LB. Phages with the broadest host range on TSB being able to infect three isolates were from the *Straboviridae* and Unclassified families; Mexia, Marfa and EI. Phages from the *Demerecviridae* were able to infect two isolates; Mineola, Matisse, Miro, Spivey, Sugarland, and Patroon. Most phages in the collection had narrow host ranges only able to infect the isolated used for propagation. No interactions beyond the propagation phages were observed strains 44809, 44818, 42195, I0201559, and I0201547.

The working culture system for phage experiments in this work has been in TSB medium, however LB medium is also commonly used for propagation and characterization of *Klebsiella* phages. The host range of most phages was found to be consistent between the two media, however the sensitivity of strains to phages in the phages in the Staraboviridae, Demerecviridae, and the myophages EI and Marfa were found to be altered on LB compared to TSB. Phages Matisse and Miro were able to plate with low efficiency on strain 39428 on LB, phage Sugarland was able to plate with low efficiency on strain 43421, and phage EI displayed the ability to plate on strain 39427 and 43421 on LB.

### 3.4 Comparison of Phage Tail Fibers/Tail Spike/Central Tail Fiber Lytic

To better understand the proteins involved in phage receptor recognition, a set of 70 proteins with potential receptor-binding functions was extracted from the 30 phage genomes in this study (Table 3). These functions were determined based on the presence of conserved domains and homology to other known phage receptor-binding proteins based on BLASTp and HHpred searches. The phages in the collection were found to express a wide variety of RBP proteins with diverse predicted arrangements, ranging from a single predicted RBP type in *Casjensviridae* and some *Autotranscriptaviridae* members, five to six predicted RBPs in the *Ackermannviridae*, and nine predicted RBP’s in the jumbo phage Miami (Table 3).

**Table 3.**
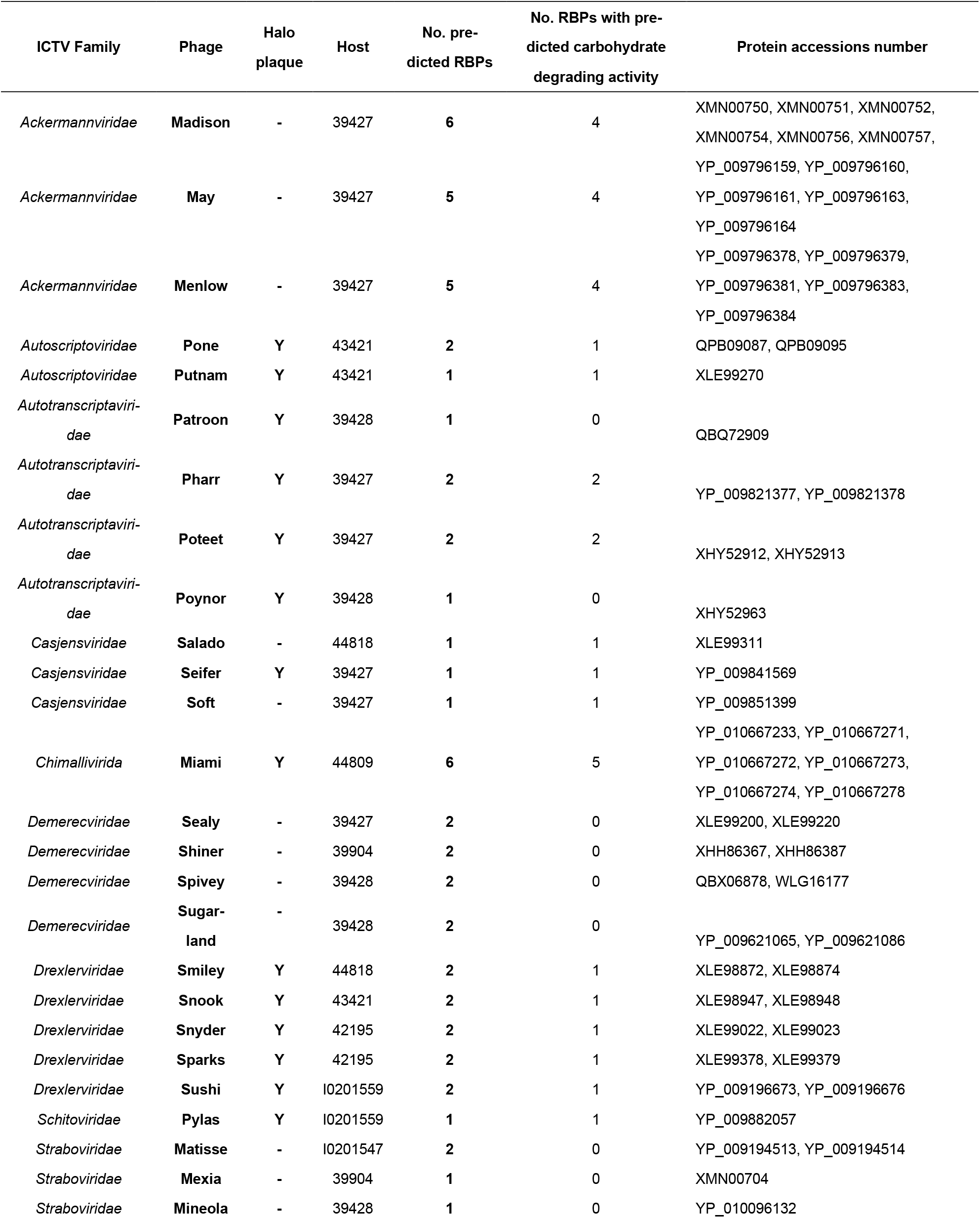
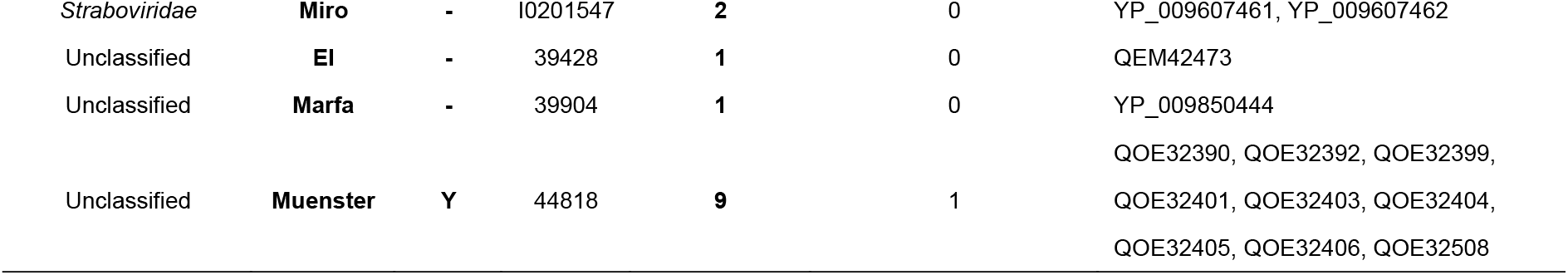
Annotated receptor-binding proteins in 30 *K. pneumoniae* phage genomes. Proteins were included based on conserved domains or similarity to other annotated phage proteins based on BLASTp or HHpred searches, 70 proteins from phages in the collection were used in the study. Phages were found to encode from 1 to 9 potential RBPs, with 17 of the 30 phages encoding at least one RBP with predicted carbohydrate-degrading activity.

An analysis of phage RBPs for potential capsular depolymerase activity identified 33 proteins across 18 phages with depolymerase-associated conserved domains or similarity to known capsular depolymerases or sugar lyases as detected by HHpred (Table 3). Phages that recognize bacterial capsule as their receptor often employ tail spikes with carbohydrate-degrading activity to adsorb to and digest their way through the capsule layer to reach the bacterial surface [66]. The presence of these depolymerase tail spikes across diverse phage families representing podo-, sipho- and myophage morphotypes demonstrates capsule may act as a potential receptor for many phage types. Some phages appear to possess multiple depolymerase tail spikes, suggesting they may be able to infect cells expressing multiple capsule types. In the majority of cases (12/18 phages), only a single depolymerase tail spike was identified (Table 3). A common indicator for the presence of a phage-borne capsular depolymerase is the presence of a characteristic turbid halo surrounding the plaque proper when plated on soft agar lawns. Based on plaque morphology (Figure 1), 15 of the 30 phages in the collection produced haloed plaques (noted in Table 3), which does not perfectly correlate with the presence of an identifiable depolymerase tail spike in the phage genome [22,26,64,66].

Previous studies on tail fiber arrangements in *Klebsiella* phages showed the presence of a wide variety of tail fiber structures in which multiple RBPs may be incorporated into a single complex branched tail fiber structure [67]. In such cases, it is often not possible to tell which of these RBPs are responsible for recognition of a given host. The predicted RBPs were grouped by phage propagation host and compared by BLASTp to identify common sequences. Both phage Smiley and Muenster infect *K. pneumoniae* strain 44818, but these phages are otherwise unrelated, as Smiley is a small T1-like *Drexlerviridae* siphophage and Muenster is an unclassified jumbo phage. However, the predicted Smiley depolymerase tail spike gp23 is clearly related to Muenster gp218, with 52% overall sequence identity that includes the predicted depolymerase domain. Given these two phages infect the same host, it appears li ely that of Muenster’s nine RBPs gp 18 is the protein responsible for host recognition. Likewise, the single depolymerase tail spike expressed by *Casjensviridae* siphophages Siefer and Soft shares 61% protein identity with *Ackermannviridae* phages May gp46, Menlow gp50 and Madison gp37. Although Madison, May and Menlow encode six, five and five RBP’s respectively, the fact that they share one RBP with Seifer/Soft which infects the same host indicates that these proteins are the ones used for host recognition in *K. pneumoniae* strain 39427.

The five *Drexlerviridae* phages in the collection appear to encode a central fiber and side tail spikes with carbohydrate degrading activity. These five phages infect four distinct hosts, with only Snyder and Sparks infecting a common host. Across these phages, the predicted central tile fiber is highly conserved (86-94% sequence identity by BLASTp), while the side tail spikes are diverse: only the Snyder and Sparks tail spikes are conserved (93% identity), while all other *Drexlerviridae* tail spikes are unrelated by BLASTp except for their extreme Ntermini which are the sites of virion attachment. These findings indicate that the side tail spikes are likely the major drivers of host specificity in these phages, and that new specificity is commonly gained via horizontal gene transfer events to acquire novel receptor-binding domains rather than by genetic drift [67–70].

## 4. Discussion

The goal of this work was to characterize a set of diverse lytic phages that are capable of infecting clinically relevant carbapenem resistant *Klebsiella* isolates, which are capable of causing a range of difficult to treat infections due to their antibiotic resistance [3,6,71]. By isolating and characterizing a diverse collection of phages against clinically relevant strains the potential of phage therapy becomes more promising alternative. In this study we characterized genomic, proteomic, and host range characteristics to reveal the diversity of phages in this collection and identified phages that could be used for potential clinical applications.

A portion of this work was to characterize the genomic features of the clinical isolates to gain better insight on the diversity of the isolate in the collection. The nine carbapenem resistant *Klebsiella* spp. isolates were selected because they were the minimally necessary strains to propagate the phages in the collection. The strains varied in sequence type (ST), with ST258 being the most represented (Table 1). ST258 is considered to be the driving force for carbapenem resistance [7,11,13]. Additionally, ST17 and ST 35 are common K. pneumoniae clonal lineages, yet not as overrepresented as ST258 [48]. While the remaining ST are not as common compared to the previously mentioned ST, including these within our collection allows for a more comprehensive of our phages against less common carbapenem resistant *Klebsiella* spp. clinical isolates.

The capsule (K) locus diversity of the strain in this collection represent a wide variety of K types; KL22, KL25, KL51, KL102, KL106, KL107, KL125, and KL142. It is necessary to have variations of K types within the collection, since capsule has been shown to be a major determinant of host tropism, it is necessary to isolate phages that can infect a variety of capsule types [22]. Since the capsule of *Klebsiella* can serve as a physical barrier to prevent phage infection, or a phage receptor [22,26]. Having strains with varied K types a more effective understanding of the host range of these phages. All strains in the collection possessed functional capsule loci, except strain 39428. This strain in the collection is a naturally acapsular, due to the truncation of a necessary capsular gene which has been shown to reduce capsule synthesis, *wcaJ* [72,73].

These phages were primarily isolated from wastewater across Texas. Genomic characterization revealed the diversity of this phage collection that spans ten ICTV families and thirteen distinct genera (Table 2, Figure 2) [34,35]. A single phage, Muenster, was unable to be classified using genome similarity of phages within ICTV. The global protein alignments within each phage family revealed high levels of gene synteny and amino acid identity within phage clades (Figure 2).

**Figure 2.**
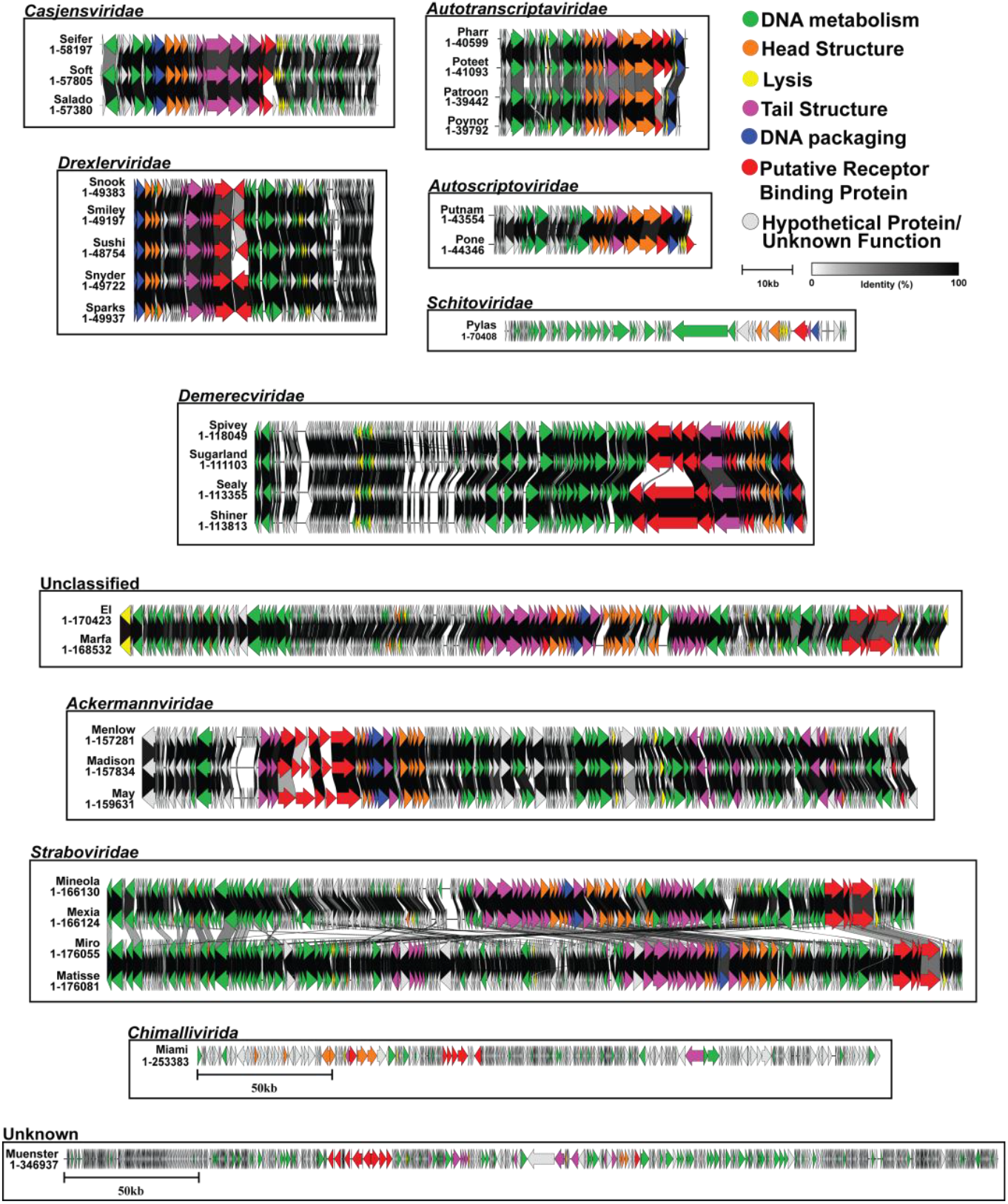
Genome similarity of isolated Klebsiella phages. The ICTV classification system was used to group and classify phages into the closest viral family and genus (if applicable). The genome organization of each phage is shown. The arrows represent protein-coding sequences and are color-coded based on functional categories. C-linker was used to perform global protein alignments and calculate similarities between clusters. Phages were named using the first letter to associate to the morphological group of the phage (M, myophage; P, podophage; S, Siphophage).

A ViPTree an alignment that generated a proteomic tree of *Klebsiella* phages, was able to provide a potential classification for phage Muenster based on protein similarity to other phages within ICTV (Figure 3) [65]. While the phages in this collection possessed high level of protein similarity to each other, they displayed large amounts of diversity to other *Klebsiella* phages. An issue was the inability to classify a phage with no genome similarity to other phages. ICTV currently uses the creation of genome-based taxa based on a core set of genes [34,35,64]. While the shift to this system better reflects the ability to more effectively reflect the genomic diversity of viruses, this issue highlights a potential draw back. By only utilizing only genomic comparisons for classifying phages by a core set of genes, phages with minimal genomic identity, yet high levels of proteomic similarity may not be classified using the current ICTV system. A previously proposed topic for a family-level criteria system was using proteome based clustering [74]. The current system leaves some phages in unclassified families,yet are grouped in a genus or left unclassified. Potentially using the current ICTV system that classifies based on a core set of genes present, then utilizing proteome clustering within genus level may better reflect the genomic relationships. Fifteen phages in the collection were observed to produce plaques surrounded by translucent zones, or “halos” when plated, indicating that these phages may utili e capsule as a receptor (Figure 1) [66]. Phages with depolymerase activity typically have a narrow host range, and are limited to infecting strains with specific capsule types [22,26]. While TSB and LB are common media, other studies conducted their experiments using LB, while this study used TSB as the standard culture medium [22,26]. Previous studies have observed the differences of achievable cell densities and phage titers comparing TSB and LB as a growth medium in *E. coli* [75]. A previous study has observed a difference in a phages plating ability in TSB compared to LB [76]. TSB differs in the nutrient content compared to LB, and the variation is dextrose. The presence of additional sugars in a medium has been shown to increase capsular polysaccharide in multi drug resistant *K. pneumoniae* [77]. We conducted a host range assay using TSB and LB to determine if there were any differences in host when on the different mediums.

**Figure 3.**
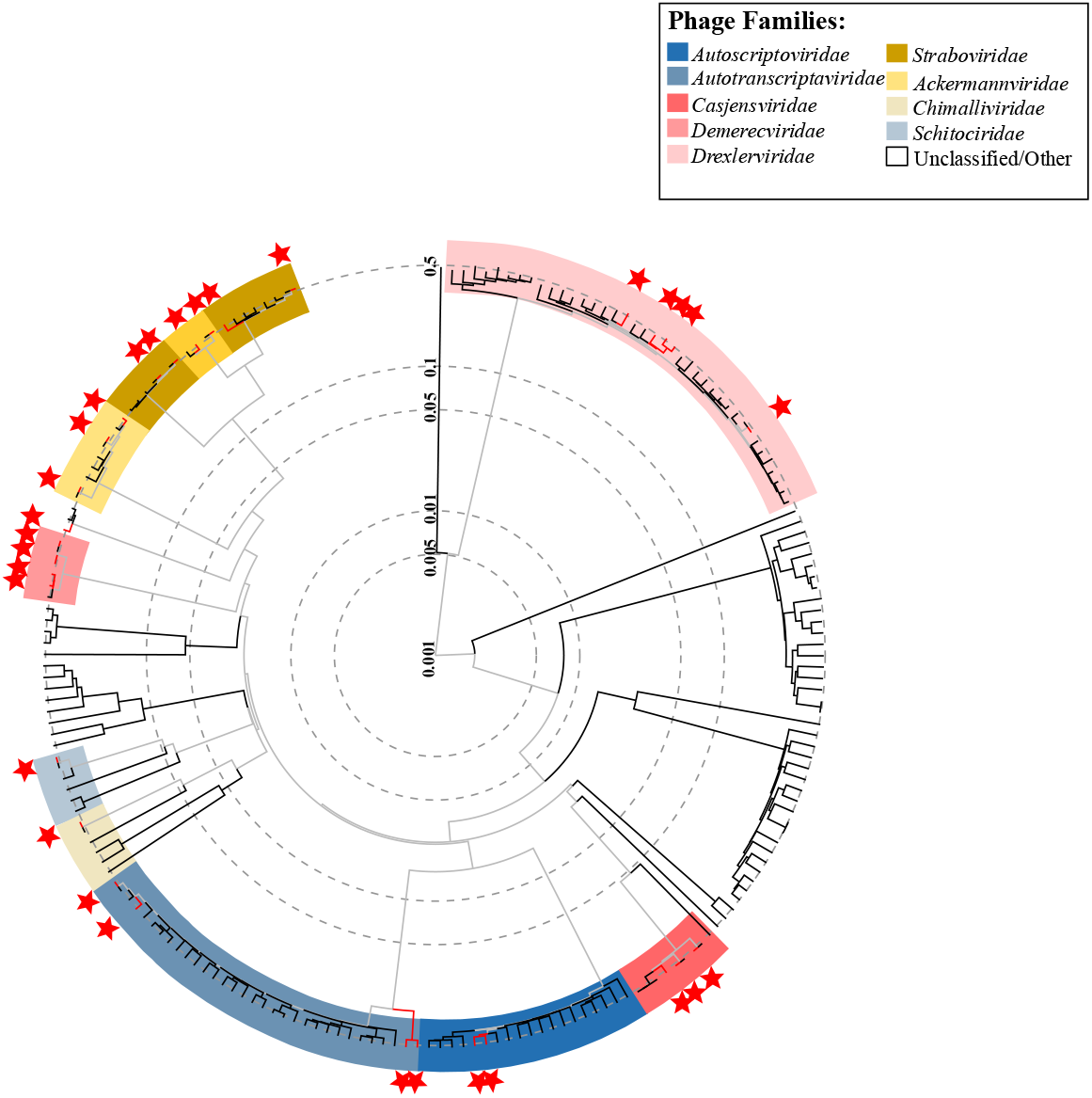
ViPTree protein-level phylogenetic tree of 30 K. pneumoniae phages. The proteomic phylogenetic tree was used to group phages based on genome wide sequence similarities. The analysis was restricted to include double stranded DNA reference phages that infect the phylum of gram-negative bacterium Pseudomonadota. This resulted in comparing 2404 of the 15,971 to the reference phages available on VIPTree against the phages in the collection to observe genome wide proteomic relationship. Phage names are colored based on ICTV Family designation.

**Figure 4.**
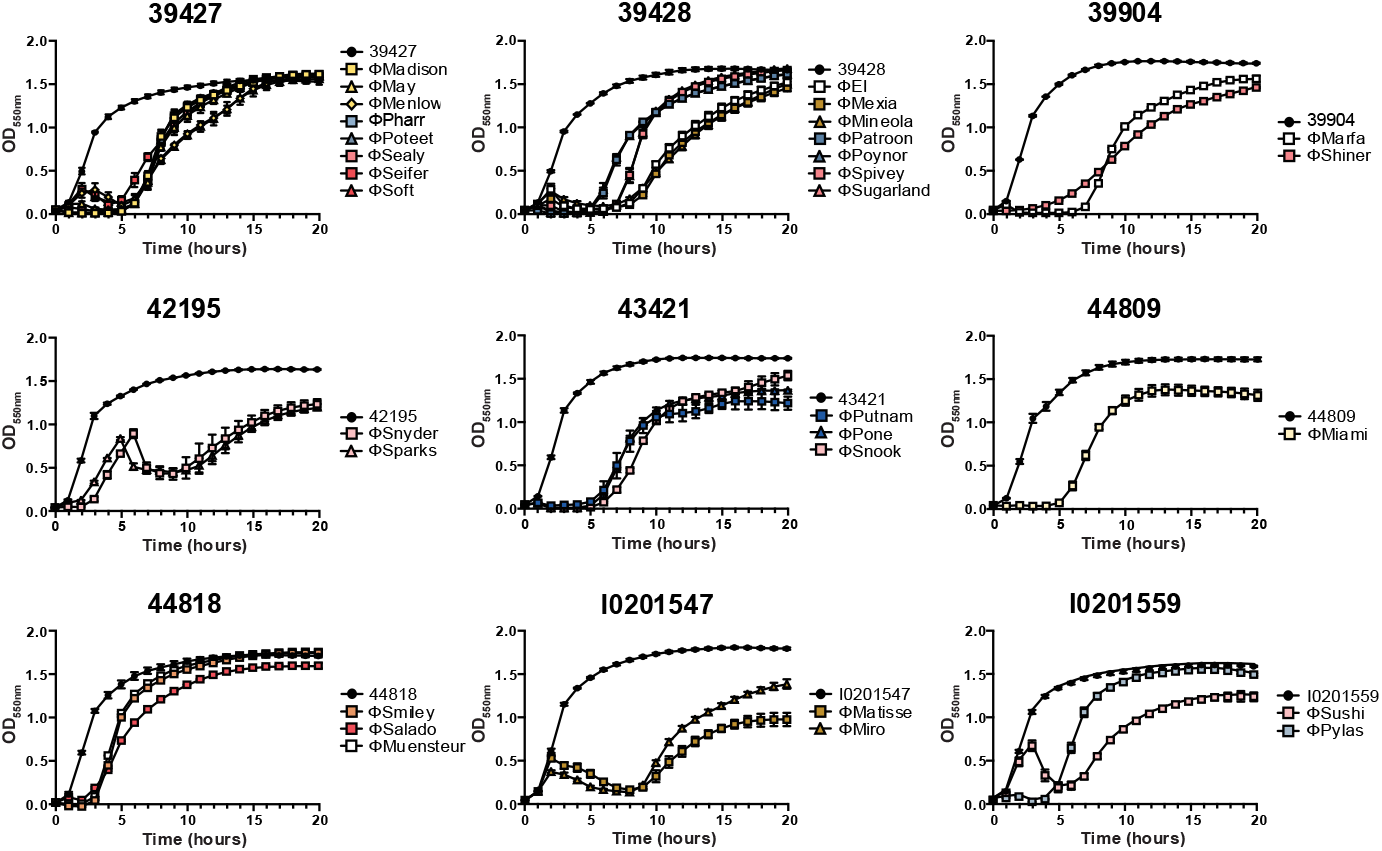
Lysis Curve. A lysis curve of each bacteriophage was generated with their respective propagation host (Table 2) at an MOI of 0.1, in TSB. The lysis curve of each phage is colored based on taxonomic Family previously using in Table 2. The black color is used for the propagation host without phage. The average of 3 independent biological replicates in triplicate are presented here.

**Figure 5.**
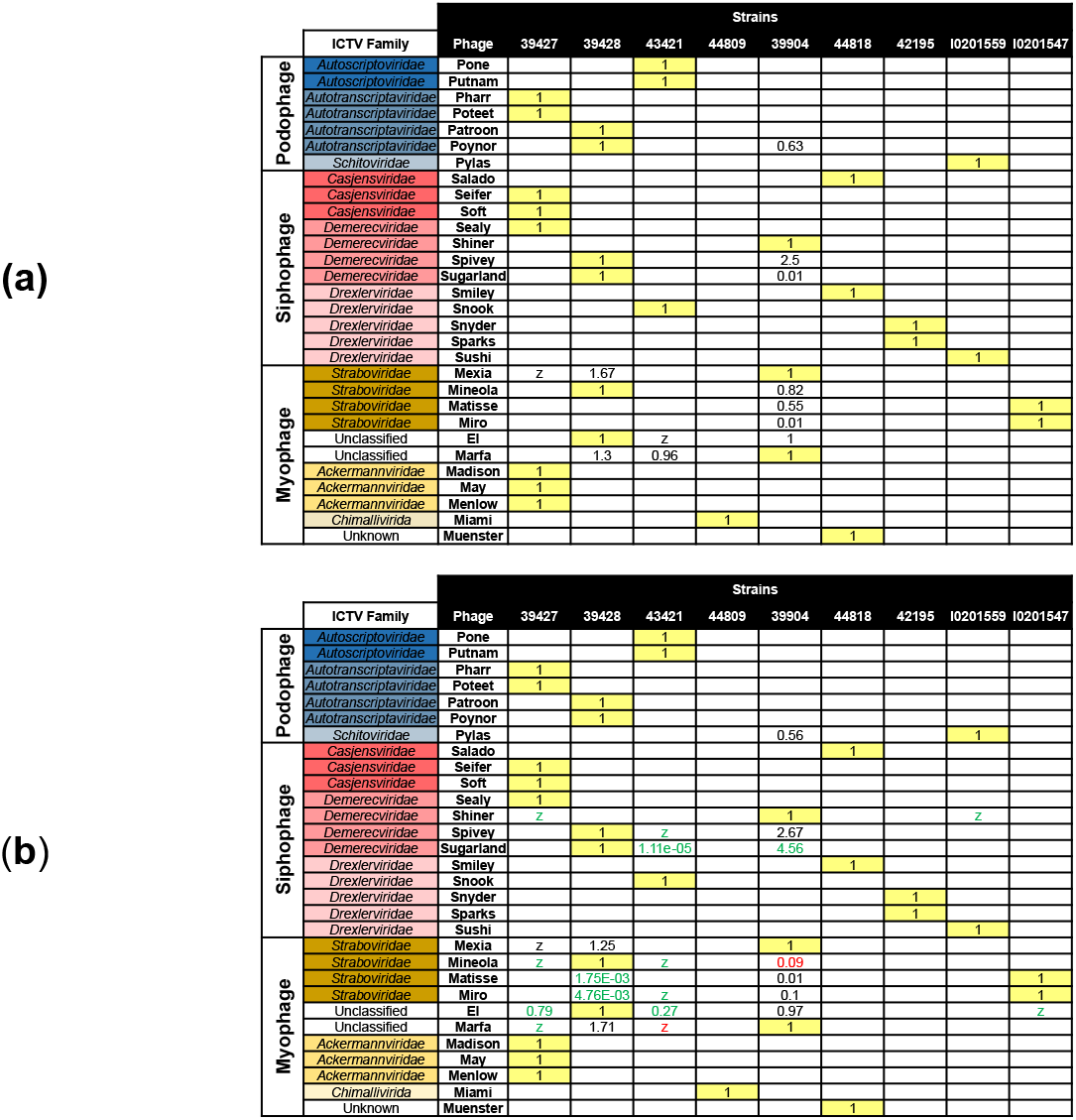
Host ranges of *K. pneumoniae* bacteriophages tested using TSB or LB culture medium. Efficiency of plating (EOP)=1 propagation host is denoted as yellow cells. A “Z” indicates the phage formed zones of clearing at high concentration by did not form plaques when further serially diluted. Blank cells indicate no interaction of the phage and strain at any concentration. Phages are separated into color groups based on their ICTV family designation, and columns are labeled with the bacterial strains tested. A: Host range of phages tested on TSB medium. B: Host range of phages tested on LB medium. Differences in host range results compared to the TSB conduction are displayed in green for an increased sensitivity and red for decreased sensitivity to the same phages; only changes of tenfold or greater are highlighted. Host ranges for most phages remain unchanged on LB, however the sensitivity to phages in the *Straboviridae* (grey), *Demerecviridae* (red) and the unclassified myophages EI and Marfa were altered on LB compared to TSB. This indicates the choice of culture medium can affect apparent phage sensitivity in host range assays.

All phages with narrow host ranges on TSB maintained narrow host ranges on LB (Figure 5). Yet the previously identified phages with a broad host range on TSB, showed an increase in host range on LB. These changes in host range were attributed to production of capsule for several reasons. Initially when bioinformatically characterizing each strain it was determined that strain 39428 is a naturally acapsular strain, due to the truncation of a necessary capsular gene which has been shown to reduce capsule synthesis, *wcaJ* [72,73]. This implicates that phages able to plate on 39428 utilize a receptor other than capsule: Spivey, Sugarland, Mexia, Mineola, Matisse, Miro, EI, Marfa, Patroon, and Poynor [22,26,64,66]. These phages were able to infect strains of various K types when plated on both TSB and LB, indicating that the type of capsule produced is not preventing phage infection but the amount of capsule. Additionally, two of these phages produce the characteristic halos and possessed proteins that indicated potential depolymerase activity; Patroon and Poynor. Yet, their ability to plate on a host that does not produce capsule further suggests that these phages do not use capsule as a receptor.

Another phage that showed an expansion of host range was Shiner. It was able to plate on strain 39427 on LB, yet not on TSB. Its propagation strain and 39427 have different K types, suggesting it may utilize a non-capsule receptor. The inability to plate on 39428 suggests potential intracellular factors or surface features preventing successful infection.

The fifteen phages that possessed halos all possessed proteins with putative depolymerase activity (Figure 1, Table 3). Several additional phages with narrow host range also encoded proteins with putative depolymerase activity: Soft, Salado, Madison, May, and Menlow. Halos are indicators for potential for depolymerase activity in a phage, which indicates that a phage may utilize capsule as a receptor. Another characteristic of phages that use capsule also possess a narrow host range and are only capable of infecting strains that possess a specific capsule type [22,26,64,66]. This may indicate that these phages may utilize capsule as a receptor. A single phage in the collection, Pharr, has a known receptor which is capsule [78].

Pharr and Poteet tail spike protein show high levels of amino acid identity, and meets other key indicators that this phage may also utilize capsule as a receptor. Phages with a broad host range showed very little to no amino acid identity with other phage receptor binding proteins. Pharr and Poteet tail spike protein show high levels of amino acid identity, indicating this phage also uses capsule as a receptor. In phages that encode multiple tail fibers, it is often not possible to determine which fiber is responsible for recognition of a given host. However, based on comparisons of tail fiber sequences it is possible in this case to assign Muenster gp218 as responsible for recognition of strain 44818 and May gp46, Menlow gp50 and Madison gp37 are responsible for recognition of strain 39427.

The findings presented here here illustrate the limitations and potential for lytic phages against CR-KP. The variability in plating on different media shows the complexity in understanding phage-host interactions, as even changes in standard media can produce different results. This work also shows our limited understanding of host receptor usage for *Klebsiella* that is not capsule. To enhance the potential utility of phages in the collection as therapeutics, is necessary to improve our understanding of the receptors used by the phages in this collection.

## Acknowledgments

The authors would like to thank Dr. Karen Frank, NIH Clinical Center, for provision of bacterial isolates. We would like to thank undergraduate student researchers Gabrielle Wilson Peterson, Madison Scott and Rachel Eckrote for assistance with experiments and genome annotations, and Lauren Lessor for technical assistance. We would also like to thank Dr. Robert Danner, NIH Clinical Center, for initial discussions and support for this project.

## Author Contributions

Conceptualization, JJG, DM, MS, TJW; methodology, DM, JJG; investigation, DM, TG, GWP, JMP; resources, JJG, TJW, MS; data curation, JJG, DM; writing—original draft preparation, DM; writing— review and editing, DM, JJG; visualization, DM, JJG; supervision, JJG; project administration, JJG, TJW; funding acquisition, JJG, TJW. All authors have read and agreed to the published version of the manuscript.

## Data Availability Statement

Phage genome and protein sequences are available from NCBI under the accession numbers provided in the manuscript.

## Abbreviations

The following abbreviations are used in this manuscript:

CR-KP: Carbapenem resistant *K. pneumoniae*
ESBL: Extended-Spectrum-Beta-Lactamase
LB: Luria Broth
LA: Luria Agar
TSB: Tryptic Soy Broth
TSA: Tryptic Soy Agar
K: Capsule
O: O-antigen
MLST: Multilocus Sequence Typing
MOI: Multiplicity of Infection
ICTV: International Committee on Taxonomy of Viruses
PFU: Plaque Forming Units
CFU: Colony Forming Units

